## Supplementary material for "DNA replication in primary hepatocytes without the six-subunit ORC": Table_1

### Embryonic lethality of *Orc2 KO*

Offspring from $Orc2^{+/\Delta}$ intercrosses

| stage | genotype $Orc2^{+/+}$ | genotype $Orc2^{+/\Delta}$ | genotype $Orc2^{\Delta/\Delta}$ | empty decidua | total # (litters) |
|---|---|---|---|---|---|
| E3.5 | 18 | 30 | 6 | 6 (n.d.) | 60 (6) |
| E7.5 | 9 | 21 | 0 | 16 | 46 (5) |
| E13.5 | 9 | 31 | 0 | 15 | 55 (6) |
| 2wk | 52 | 115 | 0 | - | 167 (27) |
