## Supplementary material for "DNA replication in primary hepatocytes without the six-subunit ORC": Table_2

Estimate of number of hepatocyte nuclei in adult mice of indicate genotypes and thus, number of hepatocyte nuclear divisions required after E9.5 mouse embryos

| | WT | *Orc2-/-* | *Orc1-/- Orc2-/- (Females)* | *Orc1-/- Orc2-/- (Males)* |
|---|---|---|---|---|
| Liver weight | 100% | 50-75% | 30% | 47% |
| Hepatocyte nuclear density | 1 | 0.5 | 0.1 | 0.66 |
| Total nuclei in liver (normalized to WT) | 100 | 25-37.5 | 3 | 31 |
| Deficit in # cell division | 0 | 1-2 | 5 | 1-2 |
| Number of nuclear divisions since E9.5 | 20 | 18 | 15 | 19 |
