## Supplementary figures and images for "DNA replication in primary hepatocytes without the six-subunit ORC"

### Supplemental_Fig_S1

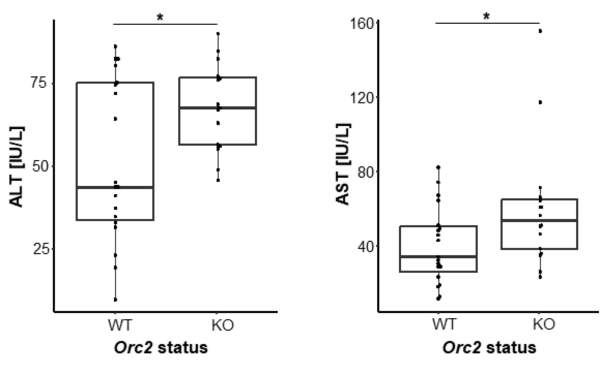

### Supplemental_Fig_S2

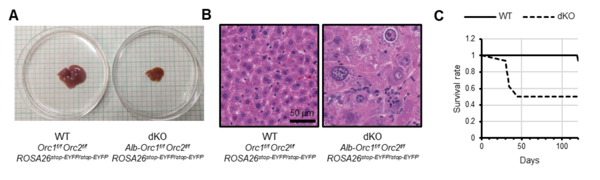

### Supplemental_Fig_S3

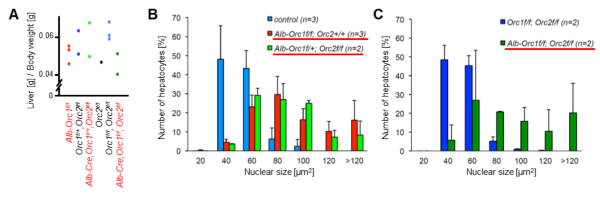
